# The hypermorphic PLCγ2 S707Y variant dysregulates microglial cell function – insight into PLCγ2 activation in brain health and disease, and opportunities for therapeutic modulation

**DOI:** 10.1101/2023.10.30.564579

**Authors:** Daniel Bull, Julie C Matte, Carmen M Navarron, Rebecca McIntyre, Paul Whiting, Matilda Katan, Fiona Ducotterd, Lorenza Magno

## Abstract

Phospholipase C-gamma 2 (PLCγ2) is highly expressed in hematopoietic and immune cells, where it is a key signalling node enabling diverse cellular functions. Within the periphery, gain-of-function (GOF) PLCγ2 variants, such as the strongly hypermorphic S707Y, cause severe immune dysregulation. The milder hypermorphic mutation PLCγ2 P522R increases longevity and confers protection in central nervous system (CNS) neurodegenerative disorders, implicating PLCγ2 as a novel therapeutic target for treating these CNS indications. Currently, nothing is known about what consequences strong PLCγ2 GOF has on CNS functionality, and more precisely on the specific biological functions of microglia. Using the PLCγ2 S707Y variant as a model of chronic activation we investigated the functional consequences of strong PLCγ2 GOF on human microglia. PLCγ2 S707Y expressing human inducible pluripotent stem cells (hiPSC)-derived microglia exhibited hypermorphic enzymatic activity under both basal and stimulated conditions, compared to PLCγ2 wild type. Despite the increase in PLCγ2 enzymatic activity, the PLCγ2 S707Y hiPSC-derived microglia display diminished functionality for key microglial processes including phagocytosis and cytokine secretion upon inflammatory challenge. RNA sequencing revealed a downregulation of genes related to innate immunity and response, providing molecular support for the phenotype observed. Our data suggests that chronic activation of PLCγ2 elicits a detrimental phenotype that is contributing to unfavourable CNS functions, and informs on the therapeutic window for targeting PLCγ2 in the CNS. Drug candidates targeting PLCγ2 will need to precisely mimic the effects of the PLCγ2 P522R variant on microglial function, but not those of the PLCγ2 S707Y variant.

**Highlights:**

- The impact of strongly hypermorphic variants of PLCγ2 have not been studied in brain and yet PLCγ2 is implicated in modifying risk of CNS disorders including Alzheimer’s disease
- To address this, we explored the role of the strongly hypermorphic PLCγ2 S707Y variant in hiPSC-derived microglia
- S707Y increases PLCγ2 enzymatic activity and intracellular calcium flux
- Phagocytosis and cytokine production are diminished in PLCγ2 S707Y microglia
- PLCγ2 S707Y downregulates expression of genes related to innate immunity and response
- Modulation of PLCγ2 for therapy must recapitulate the positive effects of moderate hypermorphic variants on microglial functions whilst avoiding detrimental effects of strongly hypermorphic variants like PLCγ2 S707Y

**Graphical abstract:** 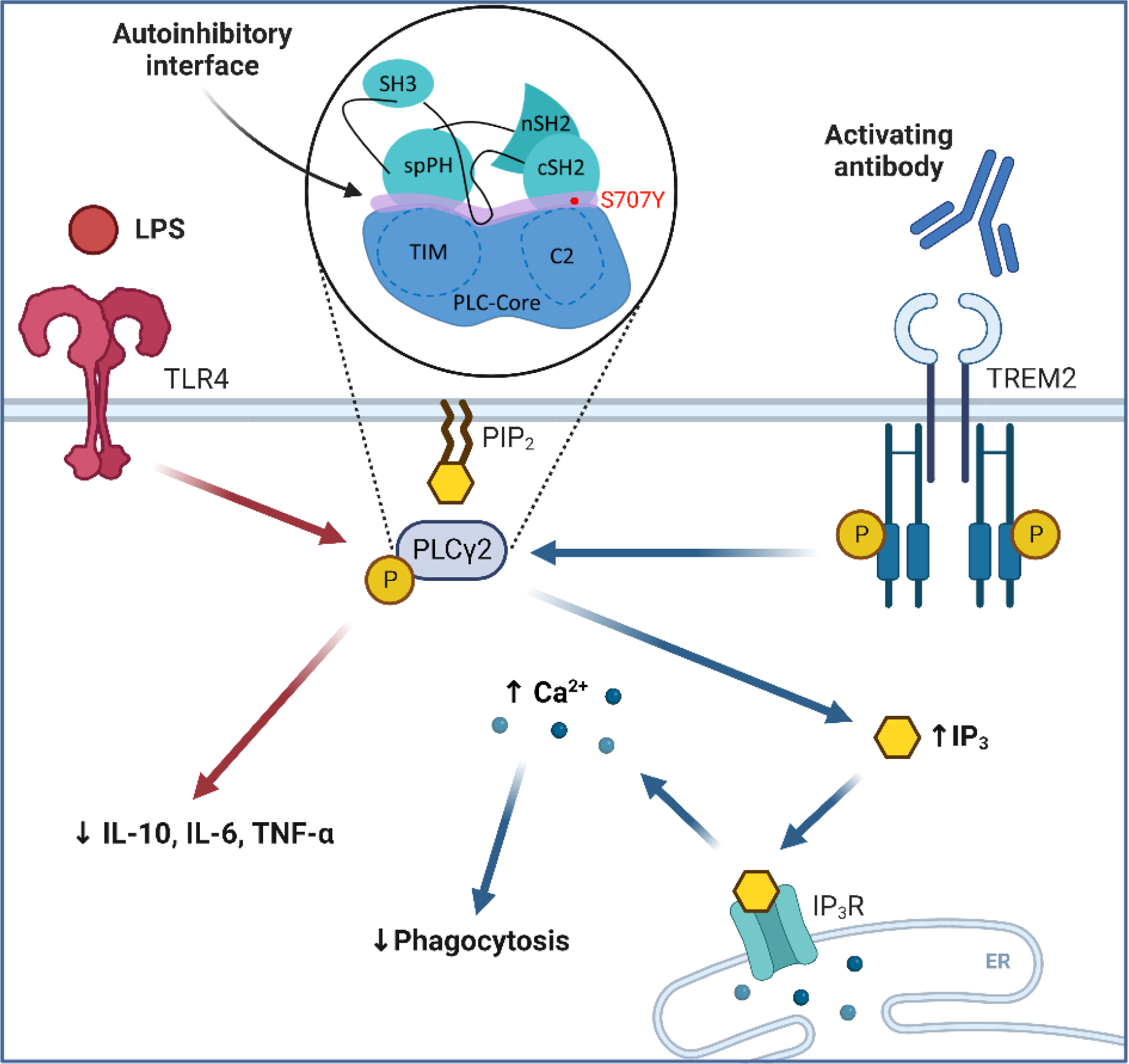

## 1. Introduction

Phospholipase C-γ2 (PLCγ2) is abundantly expressed in immune cells within the periphery and central nervous system (CNS), including microglia [1]. When activating signals are absent, PLCγ2 exists in an autoinhibitory conformation. Upon activation of specific membrane receptors or Rac GTPases, PLCγ2 undergoes a conformational change allowing it to be recruited to the plasma membrane where it catalyses the hydrolysis of the phosphatidylinositol 4,5-bisphosphate (PIP_2_) substrate into the secondary messengers; inositol 1,4,5-trisphosphate (IP_3_) and diacylglycerol (DAG) [1]. Both products propagate a wide range of downstream signals that are cell-type specific.

Human genetic studies have identified many variants of PLCγ2 that are associated with rare immune disorders, inflammatory bowel disease, drug resistance in cancer and late onset Alzheimer’s disease (LOAD) [2–6]. According to the current structural model of PLCγ enzymes, interactions between the PLCγ TIM-barrel and split pleckstrin homology (spPH) domains, as well as the C2 and C-terminal src-homology 2 (cSH2) domains, results in autoinhibition of PLCγ [7,8]. Variants with mutations that reside in the vicinity of either of these two contact points, also known as the autoinhibitory interface, can overcome enzymatic autoinhibition, resulting in increased enzymatic activity and thus alterations to downstream signalling, with one such example being the PLCγ2 S707Y (rs397514562) [3]. This variant has been identified in patients experiencing severe autoinflammatory symptoms and immune dysregulation. The term APLAID (autoinflammation and PLCγ2-associated antibody deficiency and immune dysregulation) was coined to categorise the genetic, clinical and functional findings of the PLCγ2 S707Y variant [3]; subsequently, this classification has been extended to other PLCγ2 variants which also confer strong hypermorphic PLCγ2 enzymatic activity and cause a spectrum of APLAID-related autoimmune phenotypes in this monogenic, dominantly inherited disorder [9–13]. Furthermore, it has recently been demonstrated that most of APLAID symptoms can be recapitulated in PLCγ2^S707Y/+^ mice [14].

The majority of these APLAID gain-of-function (GOF) PLCγ2 variants have been characterised in the context of immune dysregulation and autoinflammation in various immune cells of the skin, bone marrow and spleen. However, the effects of these variants on core microglial functions and phenotypes, and the possible consequences on CNS function, are only just emerging in the literature [15,19] despite recent evidence associating the rare PLCγ2 P522R (rs72824905) variant with protection against late onset Alzherimer’s disease (LOAD) and supporting healthy ageing [6,15–18]. Current functional characterisation of PLCγ2 P522R demonstrates it to be mildly hypermorphic upon activation [15,19]. In mouse models of of amyloid pathology, PLCγ2 P522R boosts microglia functions supporting containment of amyloid plaques (a key hallmark of the disease), abrogating synaptic loss, as well as rescuing working memory [16,18]. On the other hand, a loss of function (LOF) M28L PLCγ2 variant associated with increased risk of LOAD shows the opposite effects by reducing microglial responses to plaques, worsening plaque load, synaptic dysfunction and memory loss [18]. PLC enzymes are potentially druggable proteins, and, in combination with the human genetic and biology data on protective variants, this positions PLCγ2 as a very attractive target for the treatment of LOAD. Although these studies have provided an initial indication for the directionality of therapeutic modulation, it is still not fully understood how much enzymatic modulation is required to promote a therapeutic effect, without causing a detrimental response [20]. Here, we investigate the impact of a strong GOF PLCγ2 variant on microglial function, and predict a putative therapeutic window for PLCγ2 enzymatic modulation. The APLAID PLCγ2 S707Y variant was introduced into human inducible pluripotent stem cells (hiPSCs) in a heterozygous (HET) and homozygous (HOMO) fashion using CRIPSR/Cas9, before being differentiated into hiPSC-derived microglia. Compared to PLCγ2 wild type (WT) hiPSC-derived microglia, the PLCγ2 S707Y hiPSC-derived microglia exhibit hypermorphic enzymatic activity under both basal and stimulated conditions. Accompanying this large increase in variant-induced enzymatic activity, we observed a reduction in phagocytosis, as well as cytokine secretion upon inflammatory challenge, supported by bulk RNA sequencing (RNAseq) data. The protective variant with mild increase in enzyme activity promoted these important microglial functions [15–18, 51] building a body of evidence that too much PLCγ2 activation conferred by strong hypermorphic variants, like S707Y, is detrimental to core physiological responses to inflammation usually observed in healthy microglia. Our work demonstrates that strong hypermorphic variants of PLCγ2 diminish key microglial processes, further emphasising the pivotal role of PLCγ2 in microglial function.

## 2. Methods

### 2.1 Generation, genotyping, karyotyping and culturing of the hiPSC lines

KOLF2 hiPSCs (Bio Sample ID: SAMEA2398402) were genetically engineered with a CRISPR/Cas9-induced homology directed repair approach to substitute single nucleotides of the PLCG2 gene (rs397514562, C > A), followed by derivation of clonal lines as described in Bruntraeger et al. [21]. Briefly, the synthetic single guide RNAs were designed (5’-CAGACUCUCAAAAUAGGCGG-3’) and combined with single-stranded donor oligonucleotides (Integrated DNA Technologies) and Alt-R S.p. Cas9 Nuclease V3 (Integrated DNA Technologies) to form a ribonucleoprotein complex, before being introduced to the KOLF2 hiPSCs through electroporation (P3 Primary Cell 4D-Nucleofector X Kit, Program CA-137, Lonza). The resulting clones were screened by multiplexed amplicon sequencing on an Illumina Mi-Seq platform with the MiSeq Reagent Kit V2 (Illumina). Clones identified with CRISPREsso [22] to be unchanged or to be carrying indels, mono or biallelic substitutions were expanded to generate a bank of hiPSC clones. hiPSCs were cultured and DNA extracted through a DNeasy Blood & Tissue Kit (Qiagen). PLCG2 locus-specific genotyping was confirmed by GVG Genetic Monitoring (Leipzig, Germany) with the methodology outlined in Dobrowolski et al., [23]. The genetic integrity of the engineered hiPSC clones was evaluated by a qPCR-based hiPSC genetic analysis kit (Stem Cell Technologies).

All work with hiPSC and the derived cell types was performed under the respective UK legislation, ethical guidelines and approval. hiPSC lines were cultured in mTeSR1 Plus medium (StemCell Technologies) on 10 µg/mL Vitronectin XF (StemCell Technologies) coated plates. RevitaCell (Gibco) would be supplemented into the medium for reviving the cells from cryopreservation.

### 2.2 Embryoid body (EB) formation, plating and maturation

EB formation was previously described in detail from Gutbier et al. [24]. Briefly, hiPSCs were seeded into AggreWell plates (StemCell Technologies) in accordance with the manufacturer’s instructions. The next day, induction was started by exchanging 75% of the mTeSR1 plus medium with mTeSR1 plus medium supplemented with 50 ng/mL recombinant human bone morphogenetic protein 4 (rhBMP4), 50 ng/mL recombinant human vascular endothelial growth factor (rhVEGF) and 20 ng/mL recombinant human stem cell factor (rhSCF) (Peprotech). This process was repeated for the following 2 days. EB’s were then harvested and plated at a density of 1 EB/cm^2^ in GFR Geltrex (Gibco) coated flasks within myeloid factory medium (Table S1). hiPSC-derived myeloid progenitors were collected from the supernatant by centrifugation.

### 2.3 Differentiation of hiPSC-derived myeloid progenitors to hiPSC-derived microglia cells

Briefly, flasks and plates were pre-coated with 10 µg/mL of human plasma fibronectin (Sigma-Aldrich) for 1h at room temperature before washing three times with water. hiPSC-derived myeloid progenitors were seeded into a pre-coated fibronectin flask at a density of 50,000 cells/cm^2^ in microglia differentiation medium (Table S2), with half the medium replaced 2 days later. On day 4, the cells were detached with Accutase (StemCell Technologies) and collected by centrifugation before being re-plated at the same density onto fibronectin coated plates in microglia differentiation medium (Table S2). The cells were differentiated for a further 3 days (7 days total) before being ready for experimentation.

### 2.4 One-step qPCR

RNA was extracted from PLCγ2 WT and S707Y hiPSC-derived microglia from three separately generated hematopoietic inductions, at different myeloid precursor differentiation ages (early, middle and late) through TRIzol (Invitrogen) separation. TaqMan primers (Table S3) were validated with 2-fold diluted concentrations of RNA. 25ng/2uL of sample RNA was added to each well, in triplicate, onto a LightCycler multiwell 384 white plate (Roche). 3uL of the qPCR detection master mix (Table S4) was added to each well before being subjected to the thermal conditions shown in Table S5, using the LightCycler 480 System (Roche) to produce cycle threshold (Ct) values. Real-time data was analysed using the 2^-ΔΔCt^ method [25]. Ct values were normalised to the average expression of the Ubiquitin C (UBC) and Actin Beta (ACTB).

### 2.5 IP_1_ (inositol monophosphate) HTRF accumulation assay

The hiPSC-derived microglia cells were re-plated (50,000 cells/cm^2^) into a 96 well plate on day 4 of differentiation. After 7 days of differentiation, microglia derived cells were stimulated with 25 µg/mL of Goat IgG (Bio-Techne, AB-108-C) or TREM2 (Bio-Techne, AF1828) for 2h. The IP_1_ protocol was performed in accordance with the manufacturer’s instructions (Cisbio). A homogenous time-resolved fluorescence (HTRF) ratio was obtained through the PHERAstar FSX (BMG Labtech). IP_1_ data was normalised to total cell number (adherent cell-by-cell analysis) captured on the IncuCyte S3 live-cell analysis system (Satorius) prior to stimulation.

### 2.6 Calcium assay

The hiPSC-derived microglia cells were re-plated into a 384 well plate on day 4 of differentiation. Calcium 6 dye (Molecular Devices) was diluted in HBSS assay buffer (Gibco, HBSS supplemented with 1.4mM MgCl_2_, 2mM CaCl_2_, 10mM HEPES) to achieve a 1X solution. After 7 days of differentiation, media was removed from the hiPSC-derived microglia cells so that only 20uL of media remains. 20uL of 1X Calcium 6 dye was added to the cells before being placed into the incubator for 2h. Baseline calcium signal was measured on the FLIPR Tetra (Molecular Devices) before cells were exposed to either HBSS assay buffer, 10 µg/mL Goat IgG or 10 µg/mL of TREM2 at set time points. Once the calcium trace had returned to the baseline, the cells were re-stimulated with 5µM Ionomycin (Invitrogen). Area under the curve (AUC) data was analysed using Screenworks (Molecular Devices). hiPSC-derived microglia experimental stimulation AUC values were normalised to ionomycin re-stimulation AUC values.

### 2.7 Preparation of pHrodo labelled SH-SY5Y cells

Protocol was previously described in detail from Hall-Roberts et al. [26]. Briefly, confluent SH-SY5Y cells (ATCC) were dislodged and pelleted in a LoBind conical tube (Eppendorf), before washing the pellet with HBSS and centrifuging again. The resulting pellet was resuspended in live cell imaging solution (Invitrogen) supplemented with 2% paraformaldehyde (PFA, Sigma) to induce apoptosis. The cells were then mixed for 10 minutes, before HBSS was added to dilute the PFA. The cells were washed twice with live cell imaging solution through centrifugation. For every million cells, 2µL of 5mg/mL pHrodo iFL Red STP Ester (Invitrogen) was added. The cells were mixed in the dark for 30 minutes before being pelleted. Finally, the cells were diluted to a concentration of 1 million cells/mL in live cell imaging solution supplemented with 5% (v/v) dimethyl sulfoxide (DMSO). The pHrodo labelled SH-SY5Y cells were stored in the dark at −20°C.

### 2.8 Phagocytosis assay

The hiPSC-derived microglia cells were re-plated into a 96 well plate on day 4 of differentiation. After 7 days of differentiation, 50,000 pHrodo labelled SH-SY5Y cells/well were added to the microglia cells before being placed into a IncuCyte S3 live-cell analysis system, with an image captured every 10 minutes for 5h, then every 1h for a further 19h (24h total). Negative controls consisted of ‘cell only’, ‘pHrodo labelled SH-SY5Y cells only’ and 1 µM (0.01% (v/v) DMSO) cytochalasin D (Invitrogen) treatment. Live cell imaging solution was added to the negative controls to ensure the total well volume was equal. Upon addition of the SH-SY5Y cells, the plate was immediately placed into the Incucyte S3 device for fluorescence acquisition overtime (four well replicates per condition, five fields of view per well). IncuCyte analysis software provided a read for the total red object integrated intensity expressed in RCU x μm^2^/image. Phagocytosis data was normalised to the total cell number (adherent cell-by-cell analysis) captured on the IncuCyte system prior to experimentation.

### 2.9 Secreted cytokine assay

The hiPSC-derived microglia cells were re-plated into a 96 well plate on day 4 of differentiation. After 7 days of microglial differentiation the cells were challenged with ± 1 ng/mL LPS (Sigma, O55:B5) for 24h, before the cell culture supernatant was removed and immediately frozen. IFN-γ, IL-1β, IL-6, IL-8, IL-10 and TNF-α cytokine concentration was quantified with the MSD V-PLEX Viral Panel 2 (human) kit (Meso Scale Discovery) in accordance with the manufactures instructions. Cytokine values were normalised to total cell number (adherent cell-by-cell analysis) imaged on the IncuCyte S3 live-cell analysis system prior to experimentation.

### 2.10 Immunocytochemistry (ICC)

Differentiated cells were fixed with 4% PFA for 20 minutes, followed by three PBS (Gibco) washes. Cells were blocked with a solution consisting of PBS supplemented with 0.1% (v/v) TritonX (Sigma) and 10% (v/v) FBS (Sigma) for 1h at room temperature. The cells were then incubated for 24h at 4°C with an anti-IBA1 rabbit antibody (1:500, Wako). Following three PBS washes, the cells were exposed to an Alexa Fluor 488 Goat anti-Rabbit IgG secondary antibody (1:500, Invitrogen) and Hoechst (1:1000, Invitrogen) for 1h in the dark. Following an additional three PBS washes, the plate was imaged on an Opera Phenix Plus high-content imaging system (Perkin Elmer). All antibodies were diluted in a solution consisting of PBS supplemented with 0.1% (v/v) TritonX and 10% (v/v) FBS.

### 2.11 WES - Western blot

The WES was performed in accordance with the manufacturer’s instructions (Protein Simple). The primary antibodies: anti-PLCγ2 mouse antibody (sc-5283, 1:50, Santa Cruz) and anti-GAPDH mouse antibody (ab8245, 1:100, Abcam) were used. Data analysis was performed in the Compass software (Protein Simple) with PLCγ2 expression normalised to the GAPDH loading control.

### 2.12 Bulk RNASeq

RNA was extracted from PLCγ2 WT and S707Y hiPSC-derived microglia from three separately generated hematopoietic inductions, at different myeloid precursor differentiation ages (early, middle and late). Libraries were prepared using the KAPA RNA HyperPrep Kit and sequenced on an Illumina HiSeq 4000 sequencer at a minimum of 25 million paired-end reads (75 bp) per sample performed by UCL Genomics (London, England). Reads were aligned with ‘spliced transcripts alignment to a reference’ (STAR) to the human genome (hg38) [27]. Mapped reads were annotated to genes from Ensembl and quantified using featureCounts [28,29]. Differential expression analysis was performed using DESeq2 package v4.1 [30]. Principal component analysis (PCA) was performed on the top 1000 most-variable genes across all samples. Genes with false discovery rate (FDR) < 0.01 and log2 fold change (log2FC) > 1 were considered significantly deregulated, with graphical representation generated via the R Bioconductor toolset [31]. For comparison purposes, the PLCγ2 deficient hiPSC-derived microglia RNASeq dataset from Andreone et al. was reanalysed and processed using the same workflow [32].

### 2.13 Statistical analysis

Results are expressed as mean ± standard deviation. All experiments consist of at least three biological replicates (biologically distinct samples), with a minimum of three technical replicates (repeated measurements of one biological sample) per condition, unless stated otherwise. Statistical methods for analysing the various data sets are indicated directly in the figure legends. Data was analysed using Graphpad Prism 9 (GraphPad Software version 9.5.0, GraphPad).

## 3. Results

### 3.1 Generation and characterisation of PLCγ2 S707Y hiPSC-derived microglia

To determine the effects of the S707Y PLCγ2 variant, we engineered a parental hiPSC Kolf2 line to express the point mutation on one or both alleles. The PLCG2 gene from each hiPSC line was genotyped to confirm the presence of the HET and HOMO PLCγ2 S707Y (Figure S1A). As hiPSCs are prone to genomic instability, each hiPSC line was assessed for karyotypic abnormalities, with none being found (Figure S1B) [33]. Differentiated PLCγ2 WT and S707Y hiPSC-derived microglia displayed expression of ionized calcium binding adaptor molecule 1 (IBA1) protein (Figure 1A) and comparable microglial markers (IBA1, TREM2, G protein-coupled receptor 34 (GPR34), purinergic receptor P2Y12 (P2RY12), spi-1 proto-oncogene (SPI1), MER proto-oncogene, tyrosine kinase (MERTK) and PLCG2) (Figure 1B). Protein expression of PLCγ2 was also assessed, showing relatively small changes in the HET line (Figure 1C).

**Figure 1:**
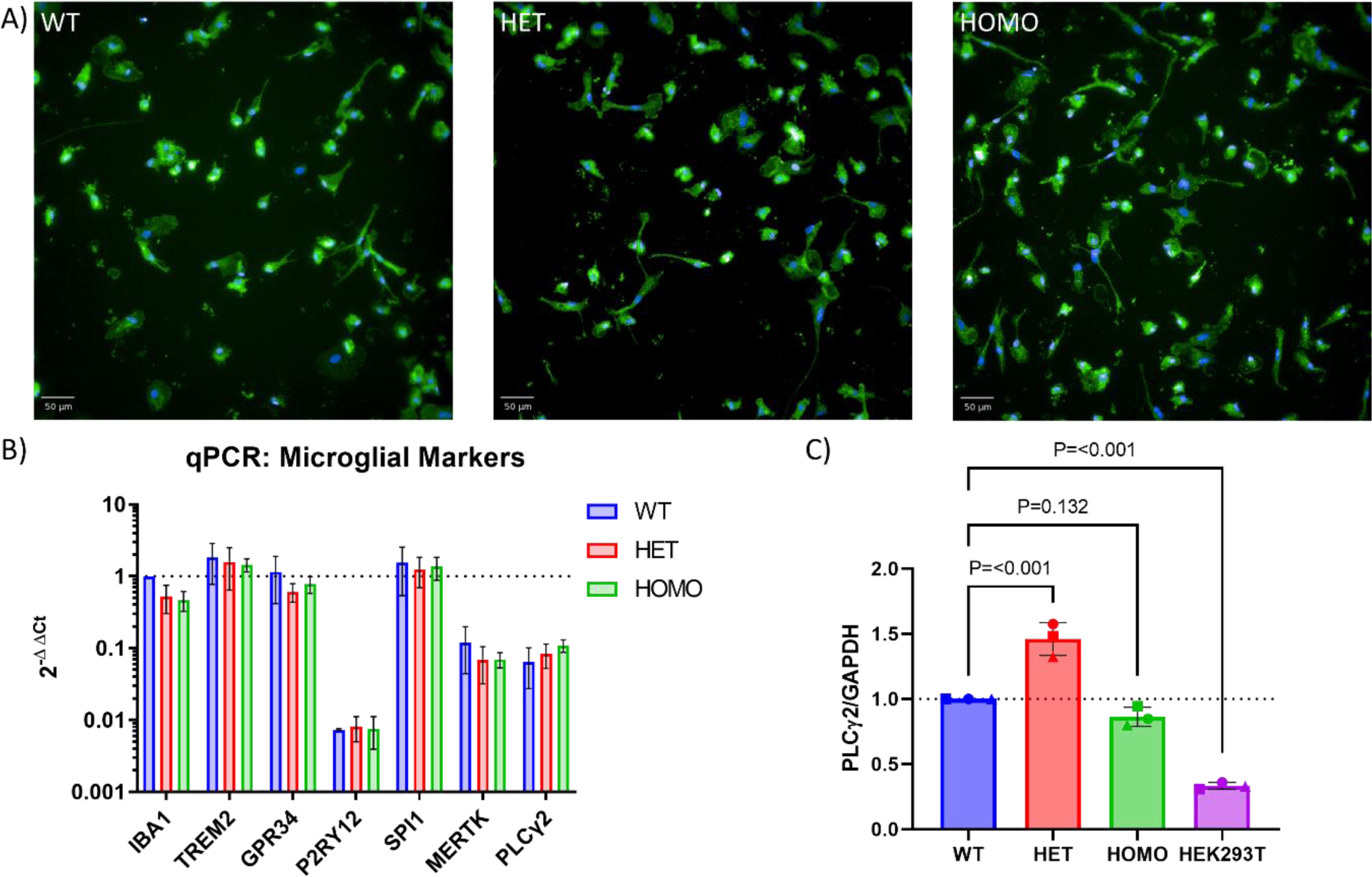
Characterisation of PLCγ2 WT and S707Y hiPSC-derived microglia. **A)** PLCγ2 WT and S707Y (HET and HOMO) hiPSC-derived microglia express calcium binding adaptor molecule 1 (IBA1). Representative images of the PLCγ2 WT and S707Y (HET and HOMO) hiPSC-derived microglia with IBA1 (green) and Hoechst (blue) staining. **B)** PLCγ2 WT and S707Y (HET and HOMO) hiPSC-derived microglia express comparable microglial markers. qPCR comparison of the difference in IBA1, TREM2, GPR34, P2RY12, SPI1, MERTK and PLCG2 gene expression between PLCγ2 WT (blue), HET S707Y (red) and HOMO S707Y (green) hiPSC-derived microglia. Ct values were normalised to the average expression of the Ubiquitin C (UBC) and Actin Beta (ACTB) housekeeping genes. Graphical values displayed were normalised to the PLCγ2 WT hiPSC-derived microglia IBA1 values from each biological replicate. Data represents the mean value ± SD from each set of biological replicates (Dunnett multiple comparisons two-way ANOVA, n=3). **C)** Bar graph displaying the relative PLCγ2 protein expression normalised to the GAPDH loading control. HEK293T cell lysate was used as a negative control to demonstrate that PLCγ2 expression is higher in the hiPSC-derived microglia compared to commonly used human cell lines. Graphical values displayed were normalised to the WT hiPSC-derived microglia expression from each biological replicate. Data represents mean value ± SD, with each graphical symbol shape (square, circle and triangle) representing each set of biological replicates from one hematopoietic induction (Multiple comparisons one way ANOVA, p-values displayed on the graph, n=3).

### 3.2 The PLCγ2 S707Y variant is strongly hypermorphic in hiPSC-derived microglia

Standard PLC functional assays using heterologous cell lines show the PLCγ2 S707Y disease-linked variant to be hypermorphic under both basal and stimulated conditions, due to the disruption that the mutation causes on the autoinhibitory interface [3,7,34]. To achieve selective activation of PLCγ2 in microglia, a commercial antibody against TREM2 was used to stimulate signalling [35,36]. The accumulation of inositol monophosphate (IP_1_), a stable downstream metabolite of IP_3_ induced by activation of the phospholipase C (PLC) cascade, was quantified using a homogeneous time resolved fluorescence (HTRF) assay. IP_1_ production was elevated in the S707Y hiPSC-derived microglia under basal (IgG, 1.3-1.4 fold increase) and TREM2 stimulated (1.1-1.2 fold increase) conditions compared to PLCγ2 WT hiPSC-derived microglia (Figure 2A, Table S6).

**Figure 2:**
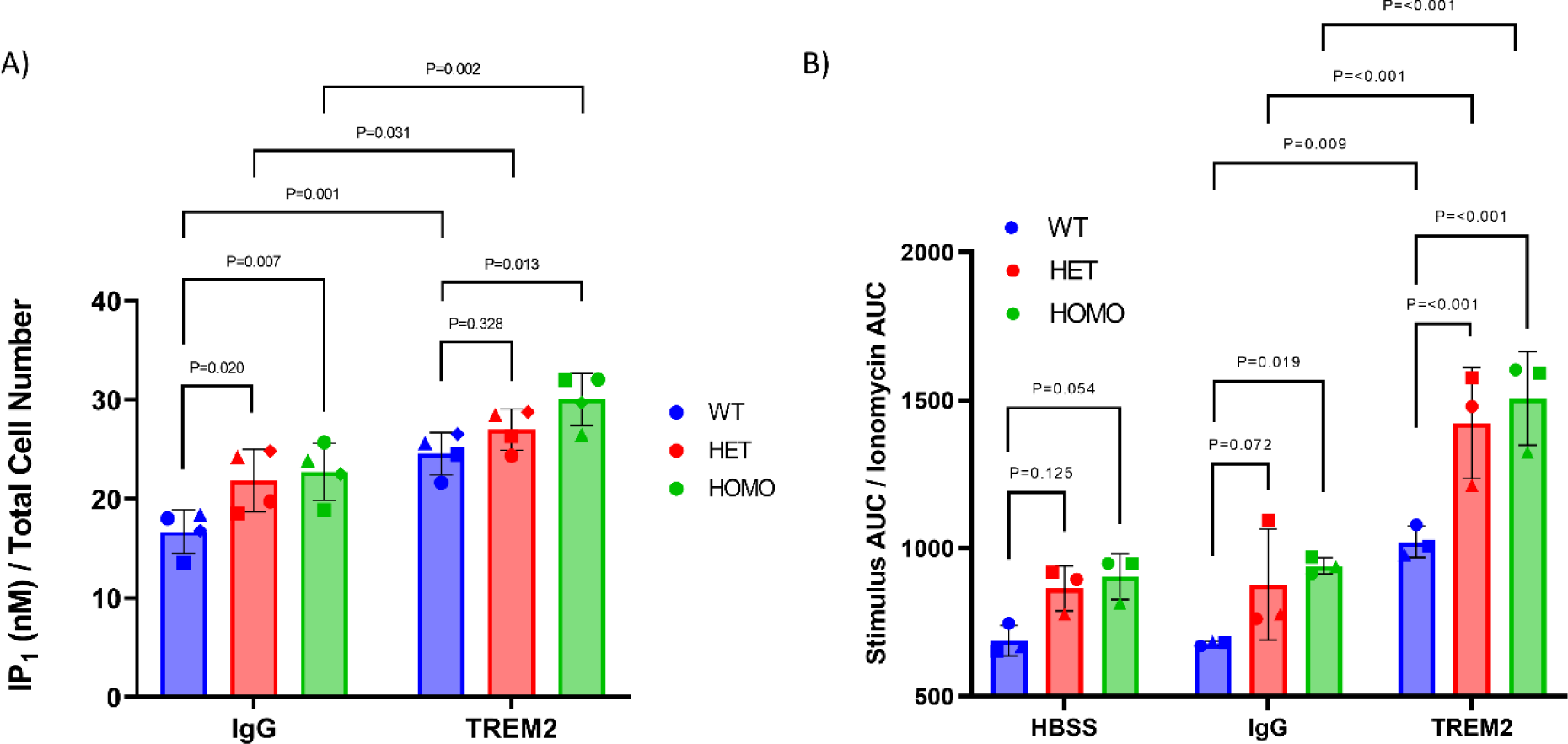
PLCγ2 S707Y hiPSC-derived microglia display increased enzymatic activity. **A)** PLCγ2 S707Y (HET and HOMO) hiPSC-derived microglia display heightened IP_1_ production compared to PLCγ2 WT hiPSC-derived microglia. Microglia were differentiated before being stimulated with 25 µg/mL of IgG or TREM2 for 2h before IP_1_ accumulation was quantified in accordance with the manufacturer’s instructions. IP_1_ values were normalised to the total cell number captured on the IncuCyte S3 live-cell analysis system prior to stimulation. Graphical data represents mean value ± SD, with each symbol shape (square, triangle, circle and diamond) representing each set of biological replicates from one hematopoietic induction (Dunnett multiple comparisons two-way ANOVA, p-values displayed on the graph, n=4). **B)** PLCγ2 S707Y (HET and HOMO) hiPSC-derived microglia display heightened calcium flux. Microglia were stimulated with either HBSS, 10 µg/mL Goat IgG or 10 µg/mL of TREM2 before the relative fluorescence units (RFU) were measured on a FLIPR Tetra. After the calcium traces had returned to baseline, each well was re-stimulated with 5 µM Ionomycin and the RFU measured. Experimental area under curve (AUC) calcium data was normalised to the AUC of the ionomycin re-stimulation data for each experimental condition. Graphical data represents mean value ± SD, with the symbol shape (triangle, square and circle) representing each set of biological replicates from one hematopoietic induction (Dunnett multiple comparisons two-way ANOVA, p-value displayed on the graph, n=3). Representative calcium stimulation kinetic traces are shown in Figure S2.

It is well documented that IP_3_ produced from PLC enzymatic function binds to the IP_3_R on the endoplasmic reticulum contributing to intracellular calcium flux (Kania et al, 2017). TREM2-stimulated PLCγ2-deficient hiPSC-derived macrophages exhibit no calcium flux, suggesting that calcium release via the TREM2 pathway is entirely PLCγ2 dependent [35]. As the magnitude of PLCγ2 mediated calcium flux is dependent on enzymatic activity, a calcium assay was utilised as an orthogonal endpoint to confirm hypermorphic activity of the S707Y microglia. Similar to the IP_1_ accumulation assay, the PLCγ2 S707Y hiPSC-derived microglia display heightened enzymatic activity under both basal (HBSS and IgG, 1.3-1.4 fold increase) and TREM2 stimulated (1.4-1.5 fold increase) conditions compared to PLCγ2 WT hiPSC-derived microglia (Figure 2B, Table S6).

### 3.3 PLCγ2 S707Y hiPSC-derived microglia exhibit reduced phagocytosis of dead neuronal cells

Intracellular calcium signalling is important for microglial functions including ramification, migration, phagocytosis and release of cytokines [38]. Although the signals that trigger phagocytosis are complex and context-dependent, microglia use phagocytosis as a way of maintaining CNS homeostasis through the clearance of debris, microbes or dead cells [39,40]. PLCγ2 signalling, mediated by TREM2 activation, has been demonstrated to be pivotal for phagocytosis [32,35]. The effect that chronic hypermorphic PLCγ2 enzymatic activity and subsequent calcium flux, as exhibited from the S707Y hiPSC-derived microglia, has on key microglial functions such as phagocytosis is not yet known.

To explore this, PLCγ2 WT and S707Y hiPSC-derived microglia were challenged with pHrodo labelled dead SH-SY5Y cells (mimicking dead neurons) for 24h with fluorescence measured kinetically (Figure 3A). The 24h total red object integrated intensity from the PLCγ2 S707Y hiPSC-derived microglia was compared to the WT hiPSC-derived microglia as a percentage change in phagocytosis (Figure 3B). Phagocytosis was completely abolished with cytochalasin D, a well-known actin polymerisation inhibitor (Figure 3B). To our surprise, a 60-70% reduction in the phagocytosis of dead neuronal cells was observed for the PLCγ2 S707Y hiPSC-derived microglia over 24h.

**Figure 3:**
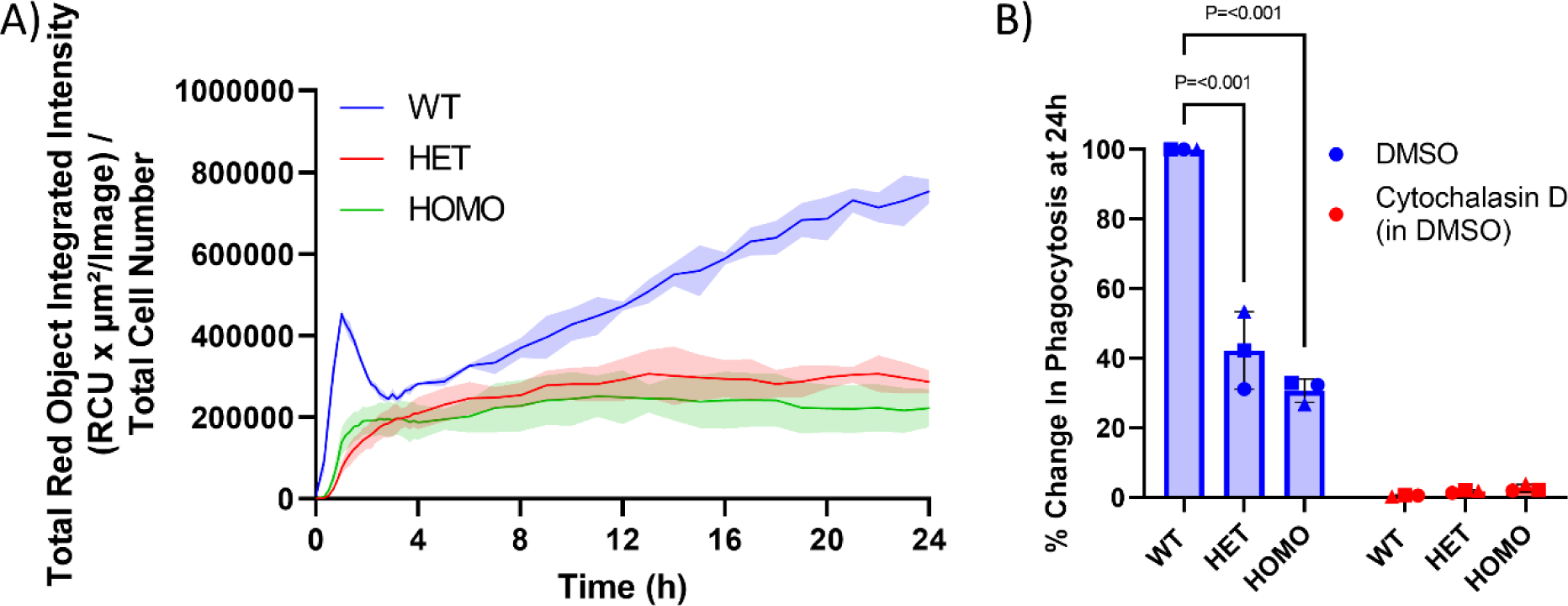
PLCγ2 S707Y hiPSC-derived microglia exhibit diminished phagocytic activity. **A)** Representative kinetic phagocytosis data of the PLCγ2 S707Y (HET and HOMO) and WT hiPSC-derived microglia. ‘No SH-SY5Y’ and ‘No microglia cell’ experimental conditions were also included as negative controls (data not shown). Images were captured on the IncuCyte S3 live-cell analysis system for 24h with the total red object integrated intensity (RCU x µm^2^/Image) analysed on the IncuCyte analysis software. Kinetic data was normalised to total cell number imaged before experimentation. Graphical data represents the mean value ± SD (Area fill). **B)** Quantification of the RCU x µm^2^/Image for the PLCγ2 WT and S707Y (HET and HOMO) hiPSC-derived microglia at 24h. Experimental PLCγ2 S707Y (HET and HOMO) RCU x µm^2^/Image values were normalised to the experimental PLCγ2 WT hiPSC-derived microglia RCU x µm^2^/Image values, and calculated as a percentage change. 1µM Cytochalasin D addition prevents phagocytic uptake (negative control). Graphical data represents mean value ± SD, with each symbol shape (triangle, square and circle) representing each set of biological replicates from one hematopoietic induction (Dunnett multiple comparisons one-way ANOVA, p-values displayed on the graph, n=3). Representative images of microglia phagocytosis at 24h are shown in Figure S3.

### 3.4 PLCγ2 S707Y hiPSC-derived microglia secrete lower levels of cytokines upon inflammatory challenge

In several reports, PLCγ2 has been shown to play a key role within TLR4 signalling [32,41,42]. Lipopolysaccharides (LPS) is commonly used to induce an inflammatory response in microglia by activating the TLR4 pathway, initiating the innate immune response, resulting in the secretion of inflammatory cytokines and chemokines [43–45]. As PLCγ2 APLAID variants, such as the S707Y, cause a strong autoinflammatory phenotype, and have shown to alter cytokine production upon LPS stimulation in peripheral immune cells, we aimed to determine whether PLCγ2 S707Y hiPSC-derived microglia would acquire an inflammatory phenotype [9,11,41].

PLCγ2 WT and S707Y hiPSC-derived microglia were stimulated with ± 1 ng/mL LPS for 24h before a panel of inflammation-associated cytokines were quantified. At the basal level, IL-8 production was increased in the PLCγ2 S707Y hiPSC-derived microglia (Figure 4). Following LPS stimulation, the PLCγ2 S707Y hiPSC-derived microglia exhibited a reduction in IL-6, IL-10 and TNF-α secretion, providing evidence that functional effects conferred by the variant are different in an inflammatory vs. basal context (Figure 4).

**Figure 4:**
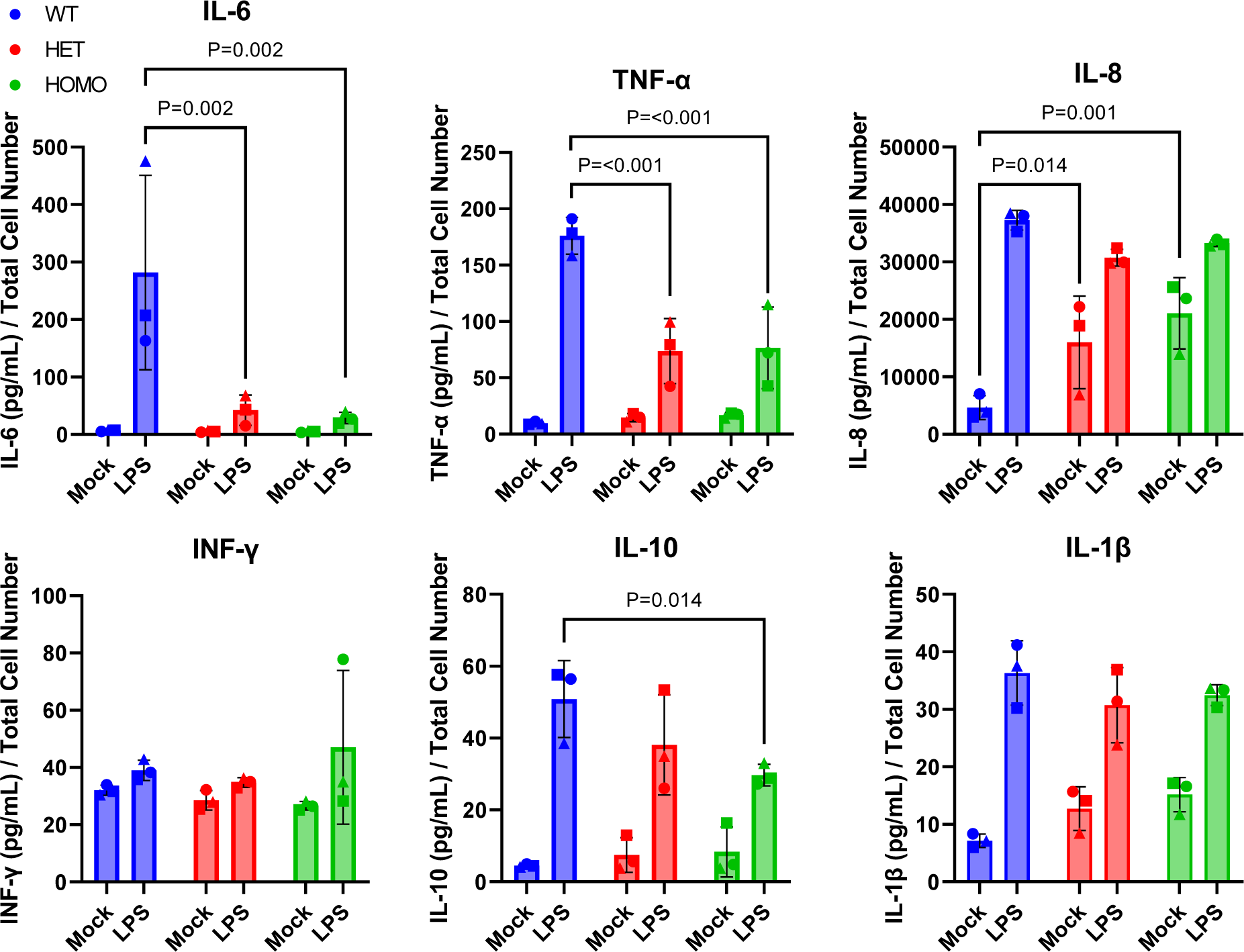
PLCγ2 S707Y hiPSC-derived microglia secrete lower levels of cytokines upon LPS challenge. PLCγ2 WT and S707Y (HET and HOMO) hiPSC-derived microglia were stimulated with ± 1 ng/mL LPS for 24h. Cell supernatant was collected from the PLCγ2 S707Y (HET and HOMO) and WT hiPSC-derived microglia, and the cytokine concentrations quantified. Cytokine values were normalised to total cell number captured on the IncuCyte S3 live-cell analysis system prior to experimentation. Graphical data represents mean value ± SD, with each symbol shape (square, triangle and circle) representing each set of biological replicates from one hematopoietic induction (Dunnett multiple comparisons two-way ANOVA, only statistically relevant p-values are displayed on the graph, n=3).

### 3.5 PLCγ2 S707Y hiPSC-derived microglia downregulate genes related to processes and responses involved in innate immunity

To investigate potential mechanisms underlying the functional effects observed, we carried out bulk RNASeq from PLCγ2 WT and S707Y hiPSC-derived microglia obtained from three separate hematopoietic inductions, and at different myeloid precursor differentiation ages, to ensure that expression changes are absolute [46]. Principal component analysis (PCA) of the top 1000 most-variable genes demonstrates separation between the PLCγ2 WT and S707Y hiPSC-derived microglia, but not between the S707Y HET and HOMO hiPSC-derived microglia, confirming similarities between the mutant lines (Figure 5A). Compared to the PLCγ2 WT, our analysis identified 766 and 701 significantly differentially expressed genes (DEGs) for the HET (Figure 5B) and HOMO (Figure 5C) PLCγ2 S707Y lines respectively, with 564 (288 upregulated and 276 downregulated) of these genes being shared amongst both PLCγ2 S707Y lines (Table S7). From the top shared DEGs identified (ranked by log2 fold change), apart from solute carrier family 15 member 4 (SLC15A4), a regulator of toll-like receptor signalling, the role of these genes in microglial function is currently unclear, and as such further deciphering of these genes is needed (Table S8) [47].

**Figure 5:**
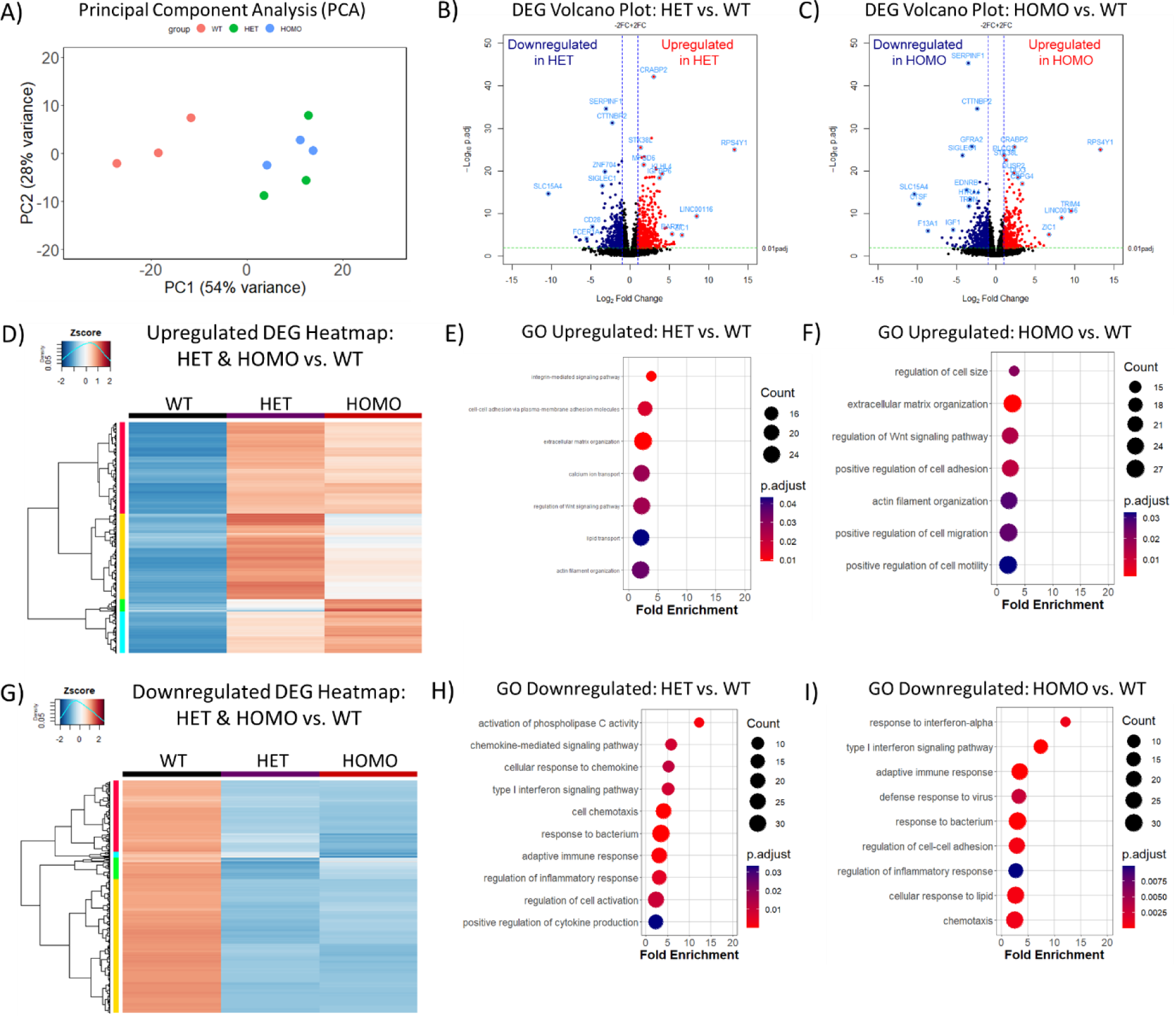
PLCγ2 S707Y hiPSC-derived microglia exhibit a downregulation of genes related to the processes and responses involved in innate immunity. **A)** Principal component analysis (PCA) of PLCγ2 WT (red), S707Y HET (green) and S707Y HOMO (blue) hiPSC-derived microglia of the top 1000 genes. **B)** Volcano plot displaying the differentially expressed genes (DEGs) downregulated (blue) and upregulated (red) in PLCγ2 S707Y HET hiPSC-derived microglia, compared to PLCγ2 WT hiPSC-derived microglia. **C)** Volcano plot displaying the DEGs downregulated (blue) and upregulated (red) in PLCγ2 S707Y HOMO hiPSC-derived microglia, compared to PLCγ2 WT hiPSC-derived microglia. **D)** Mean hierarchical cluster heatmap of the upregulated DEGs for the PLCγ2 S707Y (HET and HOMO) hiPSC-derived microglia, compared to PLCγ2 WT hiPSC-derived microglia. **E)** Gene ontology (GO) analysis of the upregulated terms in the PLCγ2 S707Y HET hiPSC-derived microglia, compared to PLCγ2 WT hiPSC-derived microglia. **F)** GO analysis of the upregulated terms in the PLCγ2 S707Y HOMO hiPSC-derived microglia, compared to PLCγ2 WT hiPSC-derived microglia. **G)** Mean hierarchical cluster heatmap of the downregulated DEGs for the PLCγ2 S707Y (HET and HOMO) hiPSC-derived microglia, compared to PLCγ2 WT hiPSC-derived microglia. **H)** GO analysis of the downregulated terms in the PLCγ2 S707Y HET hiPSC-derived microglia, compared to PLCγ2 WT hiPSC-derived microglia. **I)** GO analysis of the downregulated terms in the PLCγ2 S707Y HOMO hiPSC-derived microglia, compared to PLCγ2 WT hiPSC-derived microglia.

To further dissect the differences in gene expression between the PLCγ2 S707Y lines compared to WT, we performed hierarchical clustering for the upregulated and downregulated genes. Heatmaps confirm that the majority of the differences are attributed to the S707Y variant, independently of being HET or HOMO (Figure 5D and 5G). Gene ontology (GO) terms for the HET upregulated genes (417) are similar to the GO terms for the HOMO upregulated genes (340), as illustrated in Figures 5E and 5F. Among these, we identify cell adhesion and extracellular structure rearrangement as a term upregulated by PLCγ2 S707Y hiPSC-derived microglia (Figure 5E, Figure 5F, Table 1, Table S9). GO analysis for the downregulated genes of both HET (349) and HOMO (361) lines show similar terms, such as the downregulation of type I interferon signalling, response to bacterium, chemotaxis, PLC activity, as well as genes related to the complement response and the disease associated microglia (DAM) phenotype (Figure 5H, Figure 5I, Table 1, Table S9), suggesting a downregulation of innate immune cell processes and responses.

**Table 1:** A list of the shared and unique differentially expressed genes (DEGs) amongst the PLCγ2 S707Y (HET and HOMO) hiPSC-derived microglia and PLCγ2-deficient hiPSC-derived microglia [32]. (-) abbreviation indicates that the gene did not pass the threshold of analysis.

| Gene Name | Gene Abbreviation | PLCy2 S707Y (HET) vs. WT |  | PLCy2 S707Y (HOMO) vs. WT |  | PLCy2 deficient vs. WT |  |
| --- | --- | --- | --- | --- | --- | --- | --- |
|  |  | log2 Fold Change | adj.Pval | log2 Fold Change | adj.Pval | log2 Fold Change | adj.Pval |
| Integrin Subunit Alpha 3 | ITGA3 | **1.14** | 2.13E-03 | **1.25** | 3.49E-05 | **2.52** | 5.80E-07 |
| Integrin Subunit Alpha 6 | ITGA6 | **1.19** | 1.26E-07 | **1.36** | 6.09E-12 | - | - |
| Interferon Induced Protein With Tetratricopeptide Repeats 1 | IFIT1 | **-2.52** | 4.90E-04 | **-2.65** | 2.36E-04 | - | - |
| Interferon Induced Protein With Tetratricopeptide Repeats 2 | IFIT2 | **-1.85** | 1.19E-04 | **-1.93** | 6.59E-05 | - | - |
| Interferon Induced Protein With Tetratricopeptide Repeats 3 | IFIT3 | **-1.31** | 7.43E-04 | **-1.44** | 1.80E-04 | - | - |
| Interferon Regulatory Factor 8 | IRF8 | **-1.20** | 1.81E-09 | **-1.13** | 2.08E-08 | - | - |
| Interferon Induced Transmembrane Protein 3 | IFITM3 | **-1.97** | 1.11E-06 | **-2.21** | 3.95E-08 | - | - |
| MX Dynamin Like GTPase 1 | MX1 | **-1.87** | 3.53E-02 | **-2.32** | 6.74E-03 | - | - |
| MX Dynamin Like GTPase 2 | MX2 | **-1.53** | 1.95E-02 | **-1.65** | 1.18E-02 | - | - |
| Growth Arrest Specific 6 | GAS6 | **-2.35** | 4.73E-12 | **-2.37** | 4.17E-12 | **1.35** | 1.95E-07 |
| AXL Receptor Tyrosine Kinase | AXL | **-1.69** | 3.37E-08 | **-2.22** | 1.21E-13 | **2.21** | 3.60E-07 |
| Pyrimidinergic Receptor P2RY6 | P2RY6 | **-1.65** | 3.76E-03 | **-2.39** | 9.33E-06 | - | - |
| Complement C1q A Chain | C1QA | **-1.89** | 6.23E-08 | **-2.03** | 5.38E-09 | - | - |
| Complement C1q B Chain | C1QB | **-1.72** | 1.05E-05 | **-1.86** | 1.64E-06 | - | - |
| Complement C1q C Chain | C1QC | **-1.91** | 8.13E-09 | **-2.07** | 4.52E-02 | - | - |
| Complement C3 | C3 | **-1.23** | 1.12E-08 | **-1.21** | 2.01E-08 | - | - |
| Caspase 1 | CASP1 | **-1.89** | 3.49E-15 | **-2.09** | 3.62E-18 | - | - |
| C-X-C Motif Chemokine Ligand 1 | CXCL1 | **-1.76** | 1.60E-07 | **-1.98** | 3.19E-09 | **-1.84** | 3.03E-07 |
| C-X-C Motif Chemokine Ligand 2 | CXCL2 | **-2.46** | 5.33E-10 | **-2.23** | 1.70E-08 | **-2.66** | 3.41E-02 |
| C-X-C Motif Chemokine Ligand 3 | CXCL3 | **-2.35** | 1.46E-10 | **-1.47** | 9.00E-05 | **-3.55** | 6.73E-04 |
| C-X-C Motif Chemokine Ligand 12 | CXCL12 | **-3.40** | 2.94E-04 | **-3.93** | 4.08E-05 | - | - |
| C-C Motif Chemokine Ligand 8 | CCL8 | **-3.88** | 5.88E-03 | **-3.12** | 2.31E-02 | **-2.73** | 1.73E-03 |
| C-C Motif Chemokine Ligand 13 | CCL13 | **-4.27** | 3.03E-04 | **-5.76** | 4.29E-05 | **-1.42** | 4.93E-02 |
| CD163 Molecule | CD163 | **-1.91** | 7.36E-10 | **-2.14** | 4.16E-12 | - | - |
| Integrin Subunit Alpha M | ITGAM | **1.59** | 9.87E-03 | **1.54** | 1.49E-03 | **-1.29** | 3.83E-06 |
| Membrane Spanning 4-Domains A6A | MS4A6A | **-1.75** | 2.27E-09 | **-2.28** | 1.28E-15 | - | - |
| Paired Immunoglobulin Like Type 2 Receptor Alpha | PILRA | **-1.43** | 5.26E-04 | **-1.28** | 2.70E-03 | **-1.31** | 1.60E-06 |
| 2'-5'-Oligoadenylate Synthetase Like | OASL | **-1.69** | 1.96E-02 | **-2.02** | 4.53E-03 | - | - |

PLCγ2-deficient hiPSC-derived macrophages and hiPSC-derived microglia have also demonstrated reduced phagocytic activity when challenged with myelin and dead SH-SY5Y cells, as well as deficiencies in cellular adhesion and chemotaxis [32,35], suggesting that there is potentially overlap with the PLCγ2 S707Y phenotype. To assess this, RNAseq data from our study was compared to that generated for PLCγ2-deficient hiPSC-derived microglia [32]. Apart from the downregulation of chemokine genes, few directional DEGs were shared amongst the PLCγ2 S707Y and PLCγ2-deficient hiPSC-derived microglia (Figure S4 and Table 1). The lack of correspondence in the transcriptomic profile between the PLCγ2 S707Y and PLCγ2-deficient hiPSC-derived microglia suggests that they only partially overlap [32,35]. It should be noted that a different microglia differentiation methodology was implemented for the PLCγ2-deficient hiPSC-derived microglia compared to this study [32].

## 4. Discussion

In this study, we demonstrate that PLCγ2 S707Y hiPSC-derived microglia display elevated enzymatic activity using IP_1_ accumulation and intracellular calcium flux endpoints, consistent with previous literature findings in other cell types [3,48]. Accompanying the increased PLCγ2 enzymatic activity, we observed a reduction in important microglial functions including decreased phagocytosis of dead neuronal cells and decreased cytokine production upon inflammatory challenge. Alongside the phenotypic changes, bulk RNASeq revealed a concomitant downregulation of the expression of genes related to innate immunity; specifically the complement pathway, phagocytic ligands and receptors, as well as type I interferon signalling.

The enhanced enzymatic activity of PLCγ2 S707Y in microglial cells, measured by quantifying the IP_1_ accumulation, is relatively low compared to the effects described for this variant in B cells and overexpressing cell lines [3,7,9,49]. There are significantly lower levels of endogenous PLCγ2 expression in both primary and hiPSC-derived microglia/macrophages compared to B cells or overexpressing cells (our unpublished data, qPCR: normalised delta-delta-Ct over primary microglia; Ramos B cells, FC 8.1; hiPSC-derived macrophages, FC 0.66; WES: expression levels normalised over GAPDH; Ramos B cells, 2.47; primary microglia, 1.4; and comparison of expression in B cells versus monocyte in Human Protein Atlas dataset https://www.proteinatlas.org/ENSG00000197943-PLCG2/immune+cell, comparison of B cells and myeloid cell expression in Extended data Fig.7a in Elmentaite et al., [50]) which could explain this. Additionally, the IP_1_ accumulation assay is highly dependent on cell density; B cells are grown in suspension, with a higher number of cells per well conferring a greater signal-window compared to adherent microglia with less cells per well. Ligand-mediated activation of the BCR and TREM2 receptor differs substantially, and the interacting proteins and the downstream signalling cascade cannot be directly compared. TREM2 antibody is the only commercially available stimulus that has been shown to specifically activate PLCγ2 [35]. Additionally, WT hiPSC-derived macrophages display a maximal 1.3-Fold increase in IP_1_ and calcium levels following TREM2 stimulation [35]. Our results are comparable to published data.

Interestingly, differences in PLCγ2 protein expression were observed for the HET S707Y hiPSC-derived microglia, despite comparable PLCγ2 mRNA levels. Similar changes in PLCγ2 protein expression were observed for the M28L variant despite comparable PLCγ2 mRNA levels, providing evidence that variants may confer differences in PLCγ2 protein stability [18]. Furthermore, drastic phenotypic differences have been reported between HET and HOMO PLCγ2 P522R hiPSC-derived microglia, providing evidence for a putative gene-dosage effect [51]. It is possible that the increased HET S707Y PLCγ2 expression pushes the phenotype closer to the HOMO S707Y PLCγ2 which could explain why minimal differences between the variant harbouring microglia were observed for the majority of assays and RNASeq data. It is currently unclear why a change in PLCγ2 expression is observed for the HET S707Y hiPSC-derived microglia and this dose response could be a future area of investigation.

Given that PLCγ2 APLAID GOF variants promote a peripheral autoinflammatory phenotype, why do microglia harbouring the PLCγ2 S707Y variant appear to have a dampened immune response despite the increase in enzymatic activity? Strong PLCγ2 GOF activity could result in local PIP_2_ depletion or feedback inhibition, potentially explaining the reduction of phagocytosis observed, or could indicate that in various inflammatory states, the prey preference of microglia is modulated [19]. This was not studied here but future experiments should investigate how the level of PLCγ2 activation modulates function in different contexts and experimental conditions. Additionally, the dampened cytokine response suggests that perhaps chronically activated PLCγ2 prevents the microglia’s ability to respond effectively (e.g. the decreased IL-6, TNF-α and IL-10 production in response to LPS stimulation). Our findings could suggest that microglial function is more sensitive to the level of PLCγ2 activation compared to some other immune cells, and that a certain degree of potentiation of enzymatic activity is sufficient to trigger negative feedback mechanisms and reduced immune responses. The discrepancy between microglial and peripheral immune cells responses might also be explained by differences in microglial machinery compared to that of peripheral myeloid cells and leukocytes. PLCγ2 could trigger different signalling cascades in microglia to those in peripheral cells, or PLCγ2 modulation could be environment- or cell-state dependent. For instance, increased IL-1β and TNF-α secretion was reported in LPS-stimulated human PBMCs harbouring APLAID variants [11,41]. However, when repeated in the hiPSC-derived microglia, no change in IL-1β secretion, as well as a decrease in TNF-α secretion was observed. Recently, granulocyte-colony stimulating factor (G-CSF) has been identified as a main driver of APLAID [14]. Further work is needed to determine if G-CSF is also elevated in APLAID microglia, and to what role this would have on the CNS function. Moreover, patients harbouring APLAID mutations experience recurrent infections, antibody deficiency and autoimmunity. These phenotypes could be explained by the near complete absence of class-switched memory B cells [3]. We have not addressed this in our work as we focused on CNS microglia.

Data from our study and previous studies provides evidence that both extremes of PLCγ2 enzymatic activity (deficient and strongly hypermorphic) cause unfavourable effects in human microglia [32]. Broadly, PLCγ2 deficiency in hiPSC-derived microglia and macrophages causes reduced phagocytosis and altered cytokine responses to TLR stimulation [32,35]. In the context of neurodegeneration, the PLCγ2 M28L LOF variant has been demonstrated to exacerbate pathogenesis by dampening amyloid phagocytosis, as well as reducing IL-1β, TNF-α and IL-4 cytokine production, making it apparent that PLCγ2 variants uniquely alter the microglial transcriptome and phenotypes [18]. Although both deficient and hyperactivated PLCγ2 impair microglial function in a similar way, our RNASeq data suggests minimal overlap in mechanism, supporting the clear need to dissect the impact of each specific variant on cellular functionality [32]. We hypothesise that the PLCγ2 P522R variant lies in the “protective” zone, whereby the mild hypermorphic activity is beneficial to immune function, and not detrimentally impacting the immune response as observed for the strong hypermorphic PLCγ2 S707Y variant and the PLCγ2-deficient hiPSC-derived microglia [32,52,53]. hiPSC-derived microglia expressing PLCγ2 P522R and knock-in PLCγ2 P522R mice both show increased phagocytic activity towards amyloid-beta 1-42 [18,19,51]. Reduced uptake and downregulation of genes associated with the phagocytosis of fungal and bacterial particles was also observed in some experimental paradigms [18,19]. These divergent findings could be due to different experimental conditions and are confirmatory of the highly balanced activities of immune regulatory pathways and the importance of including experimental controls in these studies. In mouse models of amyloid pathology, the protective P522R PLCγ2 variant attenuates AD pathogenesis by enhancing amyloid uptake, reducing pro-inflammatory cytokine production, and preserving synaptic health [16,18]. These rodent studies confirm the crucial importance of PLCγ2 activity modulation in microglia for both healthy brain function and neuroprotection in *in vivo* systems, where brain networks are intact, and are additive to the cellular studies.

Like all brain cell monoculture models, hiPSC-derived microglia fail to fully recapitulate microglial biology due to the lack of interaction with other brain cell types, and rodent models do not exhibit all features of LOAD. Perhaps more complex models could provide additional insights into how PLCγ2 variants impact brain function and development e.g. synaptic pruning, as well as LOAD [54]. Moreover, due to the rarity of PLCγ2 APLAID GOF variants in the human population, and potential confounding factors due to major peripheral effects, the cognitive phenotypes of patients harbouring these variants to our knowledge has not been documented [52]. Perhaps the drastic downregulation of genes involved in the complement pathway, phagocytosis and type I interferon signalling we have observed could mean that the PLCγ2 S707Y hiPSC-derived microglia are protective against synapse loss by preventing excessive microglial synaptic pruning and pro-inflammatory responses [55,56]. Conversely, this dampened functional phenotype could also be detrimental as it could lead to ineffective toxic protein aggregate clearance, as well as dysfunctional brain development. Future work should address how PLCγ2 APLAID GOF variants influence brain development and homeostasis, as well as behaviour.

## 5. Conclusion

The hypermorphic PLCγ2 S707Y variant causes a complex impairment of immune functions, and downregulation of genes related to innate immunity in human microglia. We propose that strong activation of PLCγ2, as mediated by this APLAID variant, can contribute to CNS immune dysfunction. Our data supports caution in developing drug candidates directed at modulating PLCγ2 function, and evidence suggests that they should replicate the effects of the PLCγ2 P522R variant on microglial function but not those of variant S707Y.

## Author Contributions

D.B., L.M., M.K. and P.W. conceived and designed the study. J.M. and R.M. generated the PLCγ2 WT and S707Y hiPSC lines. D.B. conducted all experiments and performed all the data analysis (apart from the RNASeq analysis). C.N. conducted the RNAseq analysis. D.B. drafted the article and co-wrote the paper with L.M., P.W., M.K. and F.D. All authors have read and agreed to the published version of the manuscript.

## Data Availability Statement

Processed RNA sequencing data is available via the National Center for Biotechnology Gene Expression Omnibus repository under accession GSE227513.

## Acknowledgments

We would like to thank UCL Genomics for the generation of the RNASeq data. We would also like to thank Life Science Technologies Corporation.

