## Supplementary Material for "The hypermorphic PLCγ2 S707Y variant dysregulates microglial cell function – insight into PLCγ2 activation in brain health and disease, and opportunities for therapeutic modulation"

**Supplementary Figures**

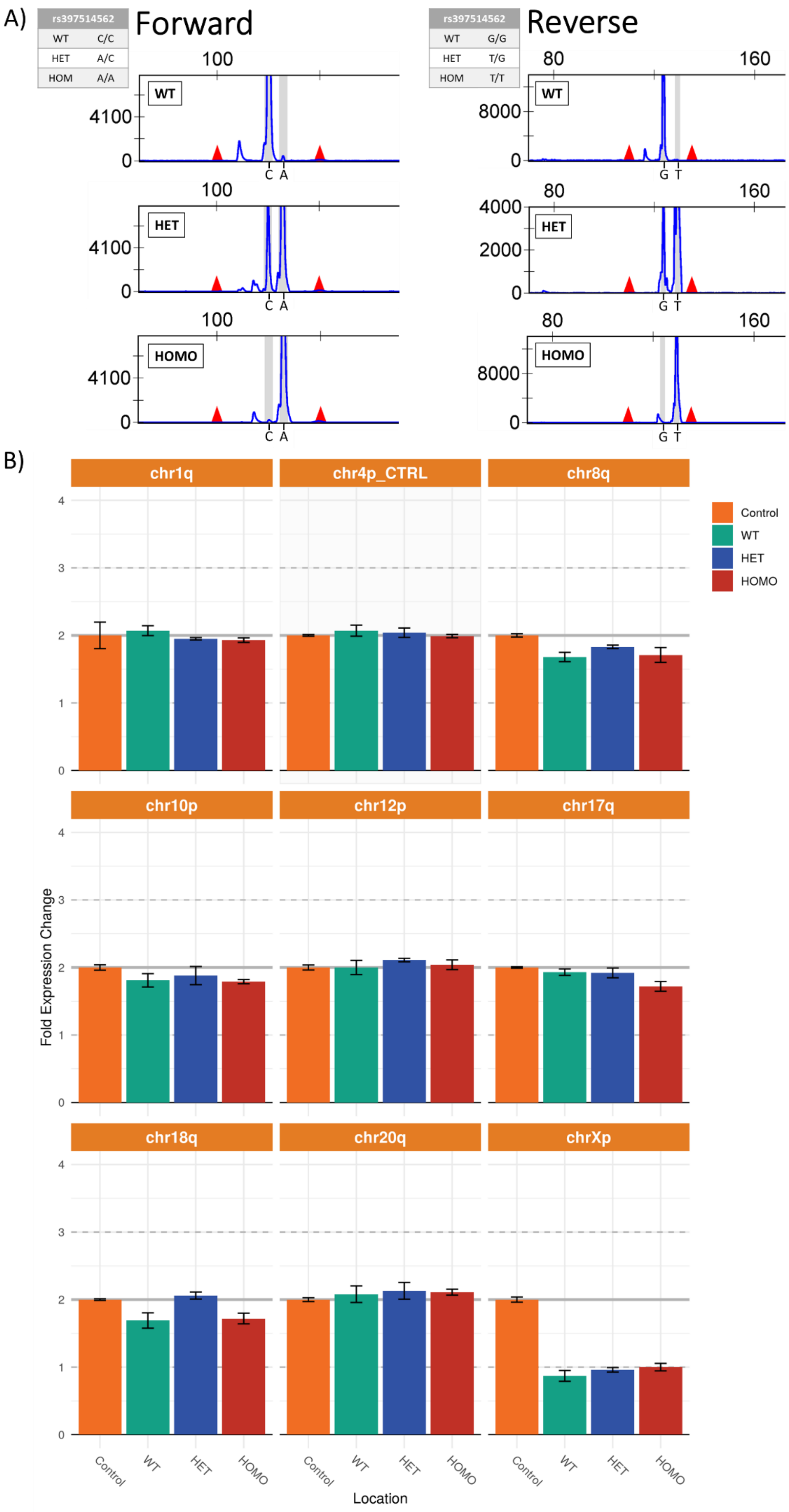

**<u>Figure S1:</u>** Karyotyping and PLCG2 genotyping of the hiPSC lines. **A)** Single nucleotide polymorphism (SNP) genotyping of the PLCγ2 WT and S707Y (HET and HOMO) hiPSC lines confirms the correct forward and reverse PLCγ2 WT and S707Y (rs397514562) nucleotides. Forward genotyping characterises between the C (WT) and A (S707Y) nucleotides, while the reverse genotyping characterises between the G (WT) and T (S707Y) nucleotides. **B)** qPCR karyotyping of the PLCγ2 WT and S707Y (HET and HOMO) hiPSC lines reveals no abnormalities in eight of the most common karyotypic abnormalities reported in hiPSCs.

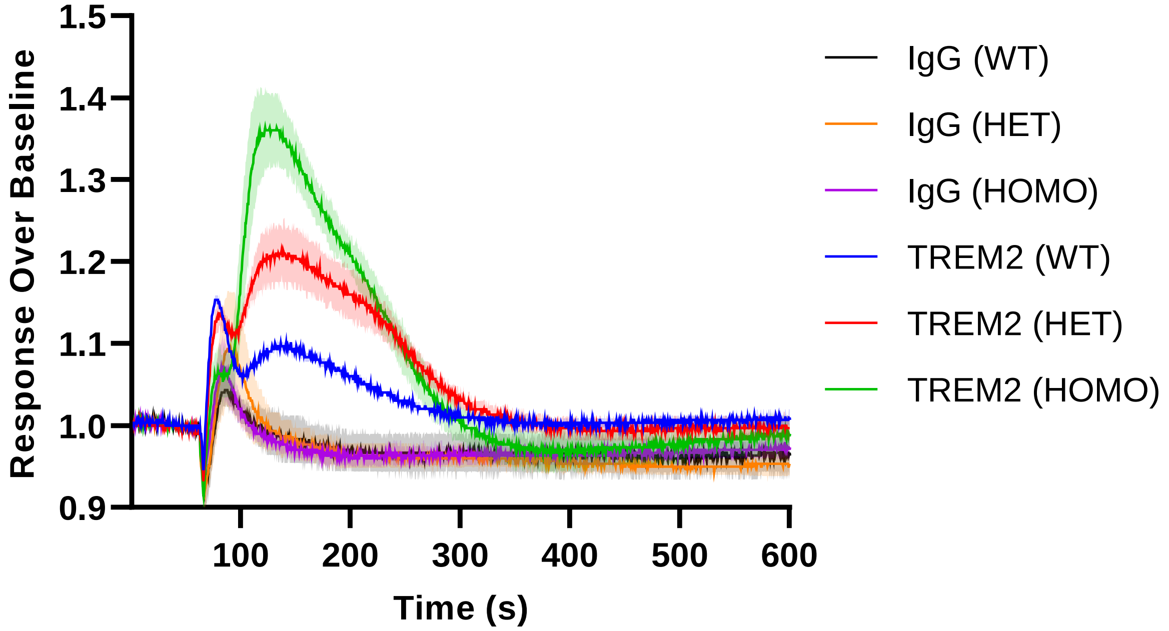


**<u>Figure S2:</u>** Representative calcium traces observed from stimulated PLCγ2 WT and S707Y (HET and HOMO) hiPSC-derived microglia. The cells were stimulated with either HBSS, 10 µg/mL control IgG or 10µg/mL of TREM2 before the relative fluorescence units (RFU) were measured on a FLIPR Tetra. Data represents mean value ± SD (Area fill).

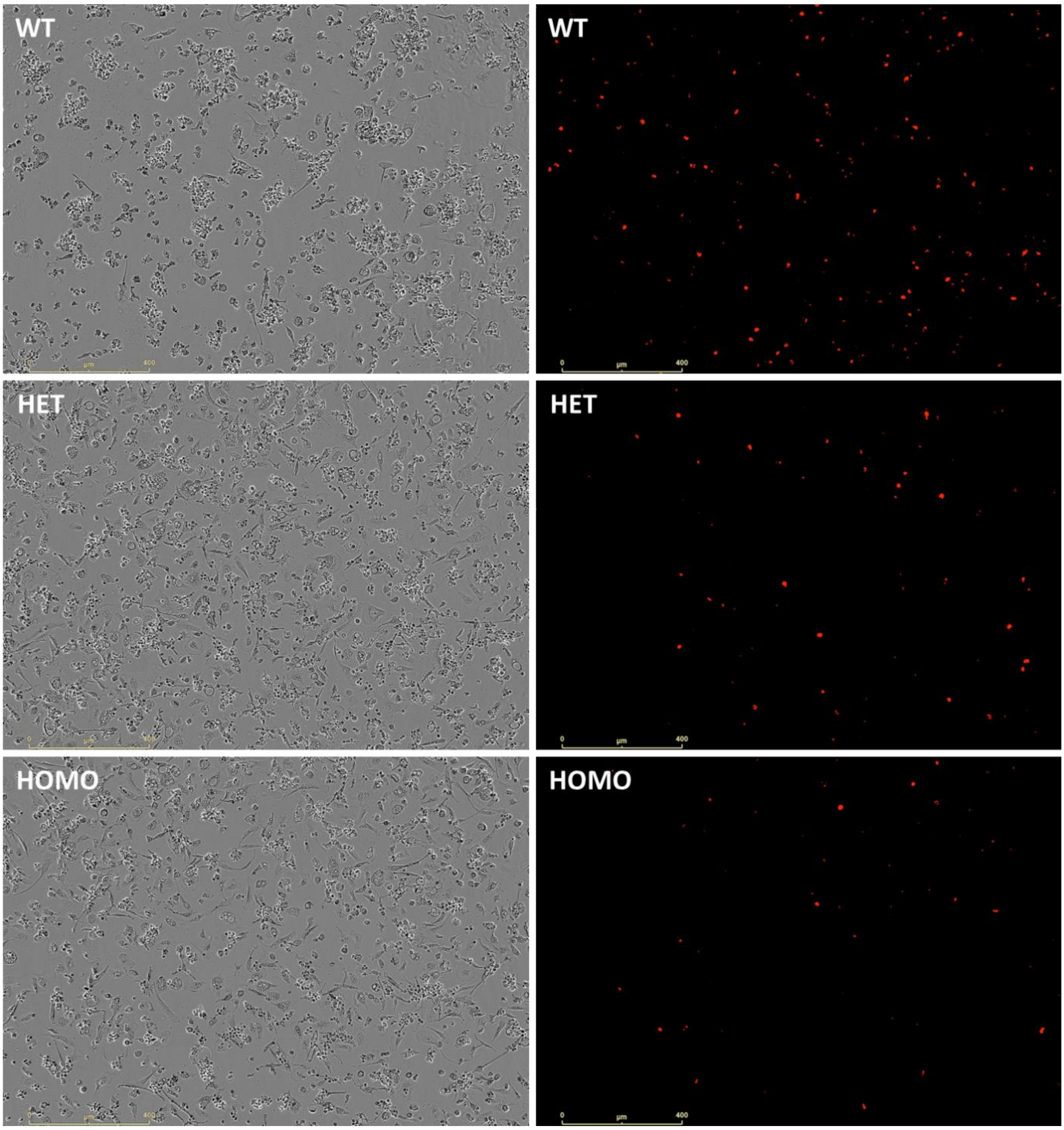


**<u>Figure S3:</u>** Representative images of PLCγ2 WT and S707Y (HET and HOMO) hiPSC-derived microglia phagocytosing pHrodo-labelled dead SH-SY5Y cells at 24h. Brightfield (phase) and red fluorescence images were captured on the IncuCyte S3 live-cell analysis system.

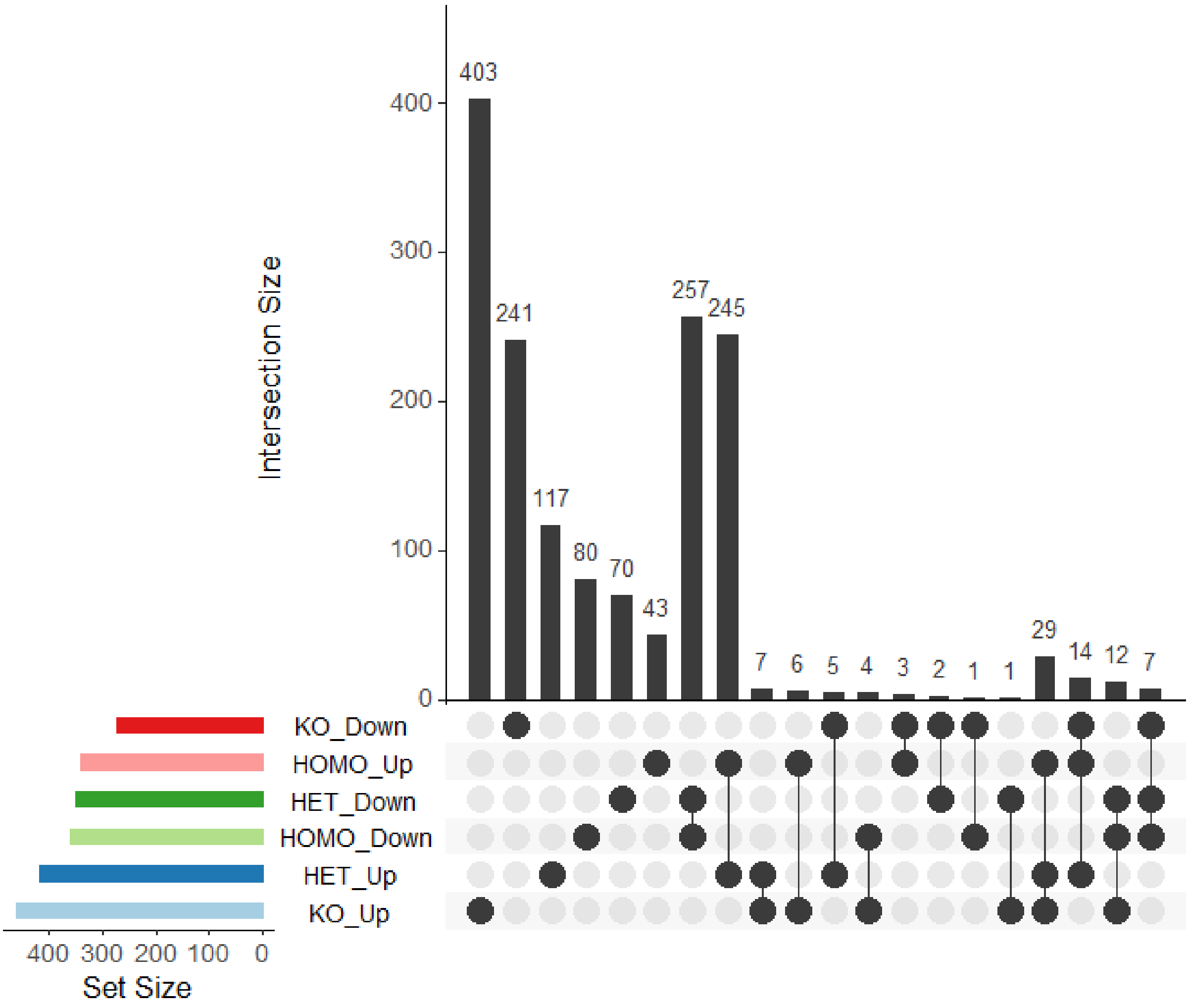


**<u>Figure S4:</u>** Visualisation of the unique and shared differentially expressed genes (DEGs) identified between the PLCγ2 S707Y (HET and HOMO) and deficient hiPSC-derived microglia [27].

**Supplementary Tables**

**Table S1:** Myeloid Factory Medium Composition

| Reagent | Catalogue Number | Manufacture | Final Concentration |
|---|---|---|---|
| X-VIVO 15 | BE02-060Q | Lonza | - |
| GlutaMAX | 35050061 | Gibco | 2 mM |
| Penicillin-Streptomycin | 15140148 | Gibco | 10 U/mL |
| β-Mercaptoethanol | 31350010 | Gibco | 50 µM |
| rhMCSF | 300-25 | Peprotech | 100 ng/mL |
| rhIL-3 | 200-03 | Peprotech | 25 ng/mL |

**Table S2:** Microglia Medium Composition

| Reagent | Catalogue Number | Manufacture | Final Concentration |
|---|---|---|---|
| Advanced DMEM/F-12 | 12634010 | Gibco | - |
| Penicillin-Streptomycin | 15140148 | Gibco | 10 U/mL |
| GlutaMAX | 35050061 | Gibco | 2 mM |
| N2 Supplement | 17502048 | Gibco | 1X |
| Cholesterol | C4951 | Sigma-Aldrich | 1.5 µg/mL |
| rhMCSF | 300-25 | Peprotech | 25 ng/mL |
| rhIL-34 | 200-34 | Peprotech | 100 ng/mL |
| rhTGF-β1 | 100-21 | Peprotech | 5 ng/mL |

**Table S3:** TaqMan (FAM) Primers

| Taqman Primer | Catalogue Number | Manufacturer |
|---|---|---|
| PLCG2 | Hs01101857_m1 | Thermo Fisher |
| P2RY12 | Hs00375457_m1 | Thermo Fisher |
| AIF1 | Hs00610419_g1 | Thermo Fisher |
| TREM2 | Hs00219132_m1 | Thermo Fisher |
| MERTK | Hs00179024_m1 | Thermo Fisher |
| SPI1 | Hs02786711_m1 | Thermo Fisher |
| GPR34 | Hs00271105_s1 | Thermo Fisher |
| ACTB | Hs01060665_g1 | Thermo Fisher |
| UBC | Hs00824723_m1 | Thermo Fisher |

**Table S4:** One-step qPCR detection master mix composition

| Reagent | Volume | Catalogue Number | Manufacturer |
|---|---|---|---|
| Water | 0.125µL | 10977035 | Invitrogen |
| FAM Primer | 0.25µL | Table S3 | Table S3 |
| RT-qPCR One step Master Mix | 2.5µL | OSPLUS-XXML | Primerdesign Ltd |
| DNAse | 0.125µL | DNASE-50 | Primerdesign Ltd |

**Table S5:** One-step qPCR thermal cycling conditions

| Step | Temperature (°C) | Duration (minutes) | Cycles |
|---|---|---|---|
| DNase Activation | 37 | 25 | 1 |
| RT | 55 | 10 | 1 |
| Taqman Hotstart | 95 | 2 | 1 |
| Denaturation | 95 | 0.1 | 40 |

**Table S6:** The percentage increase in $IP_1$ accumulation and calcium flux readouts observed for the PLCγ2 S707Y (HET and HOMO) hiPSC-derived microglia, compared to the PLCγ2 WT hiPSC-derived microglia. (-) abbreviation indicates that this stimulus was not performed.

| Assay | PLCγ2 S707Y (HET) vs. WT | | | PLCγ2 S707Y (HOMO) vs. WT | | |
|---|---|---|---|---|---|---|
| | HBSS | IgG | TREM2 | HBSS | IgG | TREM2 |
| $IP_1$ Accumulation | - | 30% | 10% | - | 36% | 21% |
| Calcium Flux | 25% | 29% | 39% | 31% | 39% | 57% |

**Table S7:**

DEG file (excel)

**Table S8:** The top five shared DEGs identified for the PLCγ2 S707Y (HET and HOMO) hiPSC-derived microglia. PLCγ2 S707Y DEGs were cross-reference with the DEGs identified from deficient hiPSC-derived microglia [27]. (-) abbreviation indicates that the gene did not pass the threshold of analysis.

| Gene Name | Gene Abbreviation | PLCγ2 S707Y (HET) vs. WT | | PLCγ2 S707Y (HOMO) vs. WT | | PLCγ2 deficient vs. WT | |
|---|---|---|---|---|---|---|---|
| | | log2 Fold Change | adj.Pval | log2 Fold Change | adj.Pval | log2 Fold Change | adj.Pval |
| Ribosomal Protein S4 Y-Linked 1 | RPS4Y1 | **13.27** | 9.79E-26 | **13.31** | 8.33E-26 | - | - |
| Solute Carrier Family 15 Member 4 | SLC15A4 | **-10.40** | 2.10E-15 | **-10.40** | 2.68E-15 | - | - |
| Mitoregulin | LINC00116 | **8.46** | 4.39E-10 | **8.35** | 9.28E-10 | - | - |
| LOC339975 | LOC339975 | **-6.89** | 3.77E-05 | **-6.90** | 4.42E-05 | - | - |
| Zic Family Member 1 | ZIC1 | **6.62** | 1.05E-05 | **6.74** | 7.95E-06 | - | - |

**Table S9:**

GO file (excel)
